# Molecular machinery of auxin synthesis, secretion, and perception in the unicellular chlorophyte alga *Chlorella sorokiniana* UTEX 1230

**DOI:** 10.1101/172833

**Authors:** Maya Khasin, Rebecca R. Cahoon, Kenneth W. Nickerson, Wayne R. Riekhof

## Abstract

Indole-3-acetic acid is a ubiquitous small molecule found in all domains of life. It is the predominant and most active auxin in seed plants, where it coordinates a variety of complex growth and development processes. The potential origin of auxin signaling in algae remains a matter of some controversy. In order to clarify the evolutionary context of algal auxin signaling, we undertook a genomic survey to assess whether auxin acts as a signaling molecule in the emerging model chlorophyte *Chlorella sorokiniana* UTEX 1230. *C. sorokiniana* produces the auxin indole-3-acetic acid (IAA), which was present in both the cell pellet and in the supernatant at a concentration of ~ 1 nM, and its genome encodes orthologs of genes related to auxin synthesis, transport, and signaling in higher plants. Candidate orthologs for the canonical AUX/IAA signaling pathway were not found; however, auxin-binding protein 1 (ABP1), an alternate auxin receptor, is present and highly conserved at essential auxin binding and zinc coordinating residues. Additionally, candidate orthologs for PIN proteins, responsible for intercellular, vectorial auxin transport in higher plants, were not found, but PILs (PIN-Like) proteins, a recently discovered family that mediates intracellular auxin transport, were identified. The distribution of auxin related gene in this unicellular chlorophyte demonstrates that a core suite of auxin signaling components was present early in the evolution of plants. Understanding the simplified auxin signaling pathways in chlorophytes will aid in understanding phytohormone signaling and crosstalk in seed plants, and in understanding the diversification and integration of developmental signals during the evolution of multicellular plants.

## Introduction

Auxin is a phytohormone that contributes to the execution of nearly all complex growth and development processes in seed plants, including gravitropism, phototropism, and cell expansion and differentiation. Indole-3-acetic acid (IAA) is the most potent and best-studied auxin in higher plants, and in *Arabidopsis thaliana* it is predominantly synthesized from tryptophan via the two-step TAA/YUC pathway: Tryptophan Transaminase of Arabidopsis 1 (TAA) converts tryptophan to indole-3-pyruvic acid (IPyA), and YUC, a family of flavin-dependent monooxygenases, converts IPyA to indole-3-acetic acid (IAA)[1]. Tryptophan-independent auxin biosynthesis has also been reported to proceed via a cytosolic indole synthase, but the enzymology of this pathway remains largely undefined [2].

Polar auxin transport proceeds through plant tissues via PIN proteins [5], whereas intracellular auxin transport between organelles proceeds through PILs (PIN-like proteins)[6]. In its protonated state, auxin can freely diffuse into the cell, where the more neutral pH cytoplasmic results in its deprotonation; therefore, the intracellular concentration of auxin is mediated primarily by efflux proteins of the ATP binding cassette family (specifically, ABCB4) and an auxin-specific amino acid permease like protein, AUX1 [7][8].

Three protein types acting in sequence mediate auxin perception in seed plants: TRANSCRIPT INHIBITOR RESPONSE 1/AUXIN SIGNALING F-BOX PROTEINS (TIR1/AFB), subunits of a SCF-type E3 ubiquitin ligase and Auxin/INDOLE ACETIC ACID (Aux/IAA) transcriptional repressors. Auxin coordinates the interaction between TIR1/AFB and Aux/IAA proteins, targeting Aux/IAA proteins for 26S proteasome mediated degradation, which in turn derepresses the transcription of auxin response factors (ARFs) (reviewed by Wang and Estelle 2014). An additional auxin sensing pathway is mediated by ABP1, a cupin domain containing protein which effects rapid, nontranscriptional auxin responses including ion fluxes at the plasma membrane [4][3].

Molecular studies of plant hormone signaling in algae have revealed a surprisingly sophisticated repertoire of plant hormone signaling orthologs [9]. The detection of auxin in the cells and tissues of charophytes and chlorophytes has been widely reported, however, the specific roles and mechanisms of auxin signaling in these algae remain obscure and a matter of dispute [10][11][12]. We now describe the molecular foundation for development of a new unicellular model for phytohormone signaling in algae. We analyzed the genome sequence of *Chlorella sorokiniana* UTEX 1230 to identify putative orthologs for auxin-active genes, and analyzed their evolutionary relationships with orthologs from other sequenced algae and plants. We thus identified candidate genes that mediate the synthesis, transport, and perception of IAA by this organism. Of particular note, *C. sorokiniana* encodes an ortholog for auxin-binding protein 1 (ABP1) in which the auxin binding residues, as well as the zinc coordinating residues essential for auxin binding, as identified in corn [13], are conserved. The functionality of this suite of genes is supported by our detection of IAA within the cell pellet and its secretion into the culture medium. This indicates a functional IAA biosynthesis and efflux pathway, with molecular mechanisms analogous to those of higher plants [14]. The observation of IAA secretion also raises the possibility that this molecule serves as an intracellular signal, allowing coordination of cellular processes across a population of cells, or within the context of a biofilm.

## Materials and Methods

### Phylogenetic analyses

We used an *in silico* approach in order to identify putative orthologs for auxin signaling in *Chlorella* and in other chlorophytes. Putative auxin signaling orthologs and outgroups were selected from representative chlorophytes, charophytes, and plants, including seed plants, ferns, and mosses. Reciprocal BLAST searches on the genome revealed candidate genes, which were aligned with Clustal Omega (except for ABP1, which was aligned with Expresso) [15] and verified by phylogenetic analysis using MEGA 6.06 using the LG+I substitution model [16]. Outgroups for these analyses were selected based on the similarity of primary protein sequences and similar domain structure, enzymatic functions, or transport functions.

*ABP1*. ABP1 was identified in the genome via reciprocal BLAST search with the *Z. mays* ortholog of ABP1 [13]. The outgroup, GLP1, was selected due to its structural similarity as a comparatively short (145 AA) protein with a cupin domain[17].

*AUX1*. AUX1 is an amino acid permease like transporter specific for auxin transport [18]. The outgroup, ANT1, is an aromatic amino acid transporter in plants [19]. Amino acid permeases tend to be highly conserved, and the substrates of AUX1 and ANT1 are of similar structure (small aromatic amino acids or amino acid-derived molecules).

*ABCB4*. ABCB4 is an ATP binding cassette transporter [7][8]. The outgroup is annotated in *Arabidopsis* as a *p*-coumaric acid transporter, and was chosen because it transports a small, aromatic molecule (as is IAA) and contains a highly conserved ATP-binding cassette domain [20].

*PILS*. PILS (PIN-like proteins) are recently discovered auxin transport proteins which are similar in structure and function, but phylogenetically distinct from the PIN polar auxin transporters in plants [6]. In order to identify the distinct lineages of PIN and PILS proteins, PILS proteins from representative plants and algae were aligned, with PIN proteins as the outgroup [5].

*IBR5*. Recently identified in *Arabidopsis* as a receptor specific for indole-3-butyric acid [21], IBR5 has orthologs that can be identified across plants and algae. MKP2 was chosen as an outgroup because both proteins contain MAP kinase phosphatase domains that are involved in complex plant hormone signaling networks [22].

### Structural modeling

CsABP1 was structurally aligned to the PDB structure of *Zea mays* ABP1 bound with napthaleneacetic acid (NAA), PDB accession 1LRH using SWISSMODEL on default settings [23]. This alignment was further investigated in PyMol v. 3.2, which was used to investigate the conservation of the auxin binding and zinc coordinating residues in CsABP1.

### IAA measurements

A *C. sorokinana* UTEX 1230 culture was obtained from the University of Texas Culture Collection of Algae. Initial cultures were grown to saturation and 50 mL cultures were inoculated at a starting cell density of 5 × 10^6^ cells/mL in Bold’s Basal Medium [24]. After two days of growth with either constant illumination or a 16h light: 8h dark photoperiod, cells were harvested. The supernatant was taken to dryness in a rotary evaporator (Büchi Rotavapor R-215, New Castle, DE) and the residue dissolved in 1 mL methanol. Cell pellets were extracted with a solution of 80% acetonitrile/1% glacial acetic acid, which was evaporated under N_2_ (N-Evap 112 Nitrogen Evaporator, Organomotion Associates Inc., Berlin, MA) and dissolved in 1 mL methanol. IAA was quantified via LC/MS/MS as described in [25]. In place of a deuterated standard, we used a standard curve of IAA ranging from 1 pM to 1 µM.

## Results and Discussion

### Conservation of tryptophan dependent IAA biosynthesis, transport, and signal transduction pathways in *Chlorella sorokiniana*

A detailed survey of the *C. sorokiniana* genome sequence revealed the presence of numerous orthologs to higher plant IAA biosynthetic enzymes, transporters, and signal transduction components, as indicated in Figure 1, with specific gene designations detailed in Table 1.

**Figure 1.**
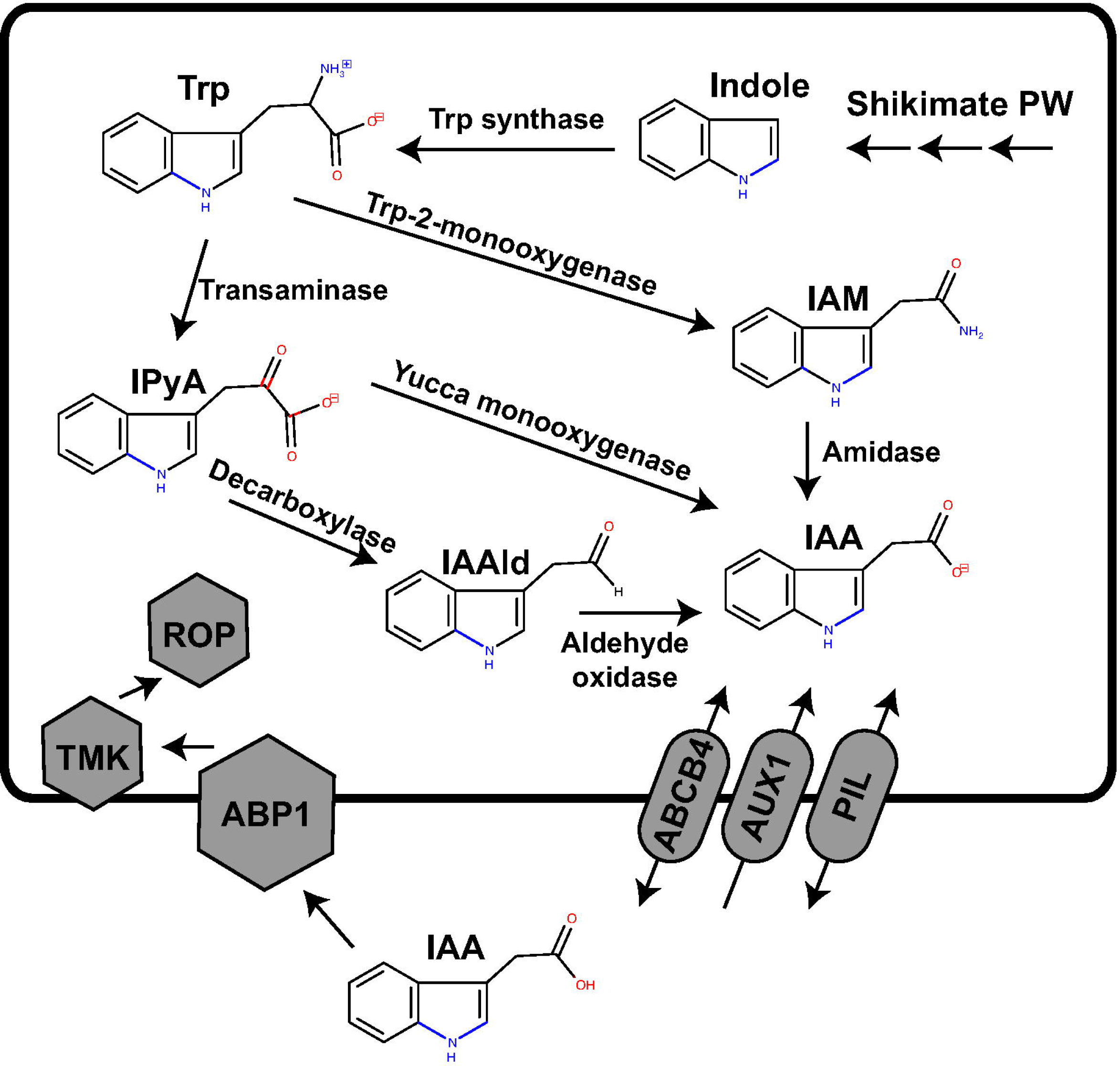
A model for auxin biosynthesis, transport, and signaling in *Chlorella*. Putative pathways of IAA biosynthesis in *C. sorokiniana* include (1.) the indole-3-pyruvic acid (IPyA) pathway: A tryptophan transaminase first converts Trp to IPyA and flavin monooxygenase (Yucca) converts IPyA to indole-3-acetic acid (IAA); (2.) The indole-3-acetamide (IAM) pathway: Primarily a pathway in phytopathogenic bacteria, IAM has been detected in all plants tested; furthermore, plants encode an amidase that converts IAM to IAA; (3.) The indole-3-acetaldehyde (IAAld) pathway. In the IAAld pathway, IPyA is converted to IAAld by a decarboxylase, and indole-3-acetaldehyde oxidase converts IAAld to IAA. Efflux and import functions are associated with the presence of specific transporters and the ABP1 receptor and associated downstream signaling components are present, as specified in Table 1.

**Table 1.** The *Chlorella sorokiniana* genome encodes putative orthologs for the biosynthesis, transport, sensing, and signal transduction of auxin. *Arabidopsis* queries were used to identify BLASTP hits in the draft *C. sorokiniana* genome. Reciprocal hits are denoted with an asterisk (*). Three newly discovered putative transaminases, not specific to tryptophan, are denoted by the diamond (^◊^).AtYUC and the putative CsYUC genes, denoted with a bullet (^●^). are not perfectly reciprocal hits, even though AtYUC queries return CsYUC and CsYUC queries against *Arabidopsis* return AtYUC among the top 10 hits.

| Functional category | Gene name | *Arabidopsis* accession | *Chlorella* accession | Percent identity | Expect | Specific function |
| --- | --- | --- | --- | --- | --- | --- |
| Receptor | ABP1 | AT4G02980 | sca021.g104100.t1* | 39 | 9e-35 | Receptor |
| Amino acid transaminases | TAA1 | AT1G70560 | sca134.g100350.t1<sup>◇</sup><br>sca110.g101700.t1<sup>◇</sup><br>sca003.g105450.t7<sup>◇</sup> | 25 | 7e-04 | *C. sorokiniana* general transaminases |
| Biosynthesis | YUC | AT2G33230 (YUC7) | sca096.g103150.t1<sup>●</sup> | 27 | 3e-14 | Indole-3-pyruvate monooxygenase |
|  | AMI1 | AT1G08980 | sca003.g101900.t1* | 39 | 3e-75 | Indole-3-acetamide hydrolase |
|  | AAO1 | AT5G20960 | sca130.g100900.t1 | 29 | 4e-119 | Indole-3-acetaldehyde oxidase |
| Transport | PILS | AT1G71090 (PILS2) | sca099.g100650.t1* | 32 | 2e-21 | PIN-like intracellular auxin transporters |
|  | ABCB4 | AT2G47000 | sca090.g105800.t1* | 39 | 0.0 | Auxin-specific ABC-type transporters |
|  | AUX1 | AT2G38120 | sca028.g105050.t1* | 34 | 1e-21 | Aromatic amino acid permease like auxin transporter |
| Signal transduction | TMK1 | AT1G66150 | sca128.g101450.t1 | 26 | 2e-35 | Activates Rho-like GTPase signaling |
|  | ROP2 | AT1G20090 | sca139.g104200.t1 | 32 | 4e-22 | Rho-like GTPase signaling |

In contrast to higher plants, a tryptophan specific transaminase was not identified with high confidence, however three putative general amino acid transaminases were identified (Table 1). This finding of a limited number of general amino acid transaminases agrees with prior knowledge that most microbial transaminases are relatively nonspecific in their activity. It also suggests that the transaminases proliferated and became more specified in their function throughout higher plant evolution. Similarly, *C. sorokiniana* encodes genes for the formation of IAA by three different routes (Fig. 1; Table 1): CsAMI1 for indole-3-acetamide hydrolase [26], CsYUC for indole-3-pyruvate monooxygenase [27], and CsAAO1 for indole-3-acetaldehyde oxidase. The amino acid sequence of CsYUC is closest to *Arabidopsis* YUC5 and YUC7, which are expressed in roots [27], though no reciprocal BLAST hit was found. *Arabidopsis* TAA does not return a transaminase specific for tryptophan: the three putative orthologs listed in Table 1 are hits for the general amino acid transaminase family, as suggested by their expect values. However, most microorganisms have a limited number of amino acid transaminases able to interact with multiple amino acids. For instance, *E. coli* has four major transaminases, each of which can interact with 3-6 amino acids [28]. Similarly, *S. cerevisiae* has four multisubstrate transaminases which participate in fusel alcohol formation via the Ehrlich pathway with two of these transaminases: Aro8p and Aro9p, neither of which have specific orthologs in *C. sorokiniana*, being broad substrate specificity transaminases for the aromatic amino acids [29]. *Arabidopsis* encodes eleven YUC family flavin containing monooxygenases whose expression is tissue dependent, reiterating the theme of proliferation and recruitment of enzymes to specific pathways and locations throughout the course of plant evolution. It does not contain any PIN orthologs.

### Conservation of IAA transporters

*Arabidopsis* contains at least four classes of auxin transporters (Figure 1, Table 1): PIN proteins, which are efflux transporters that mediate polar auxin transport [5]; PILS (PIN-like) transporters, which control intracellular auxin gradients [6]; ATP binding cassette (ABCB) transporters whose directionality depends on the concentration of auxin [7], and AUX1/Like-AUX1 (LAX) transporters [18], which are plasma membrane-localized auxin permeases that resemble aromatic amino acid permeases. *C. sorokiniana* encodes at least one putative ortholog for three of the four families of transporters: PILS, AUX1, and ABCB type transporters (Figures 2–4, Table 1).

**Figure 2.**
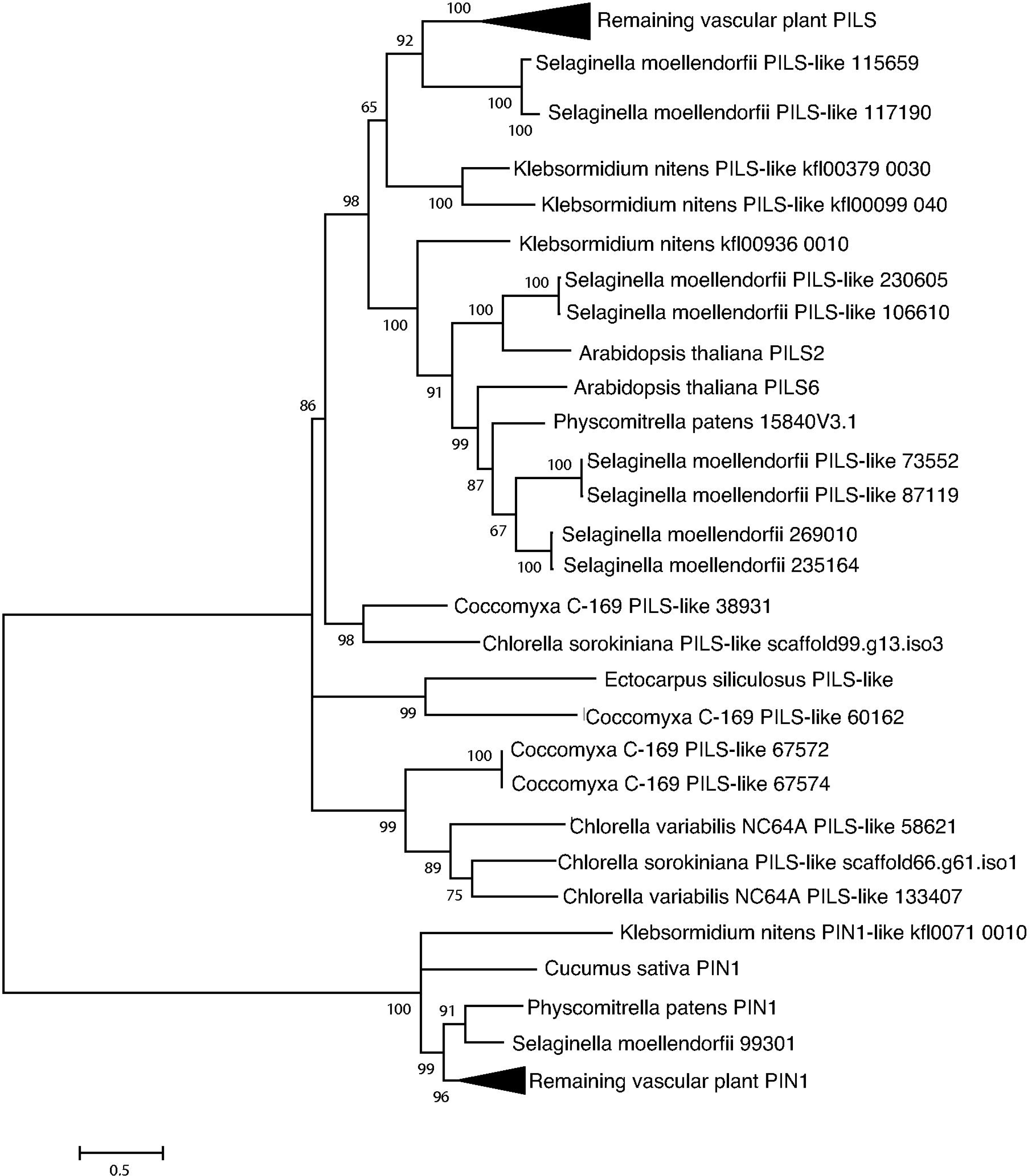
Phylogenetic analysis indicates sequence conservation of PILS. PILS (PIN-like) transporters, similar to but distinct from PIN proteins, mediate intracellular auxin transport. Putative orthologs and outgroups were identified by reciprocal BLAST searches, aligned with Clustal Omega and analyzed using the LG+I substitution model in MEGA 6.06.

**Figure 3.**
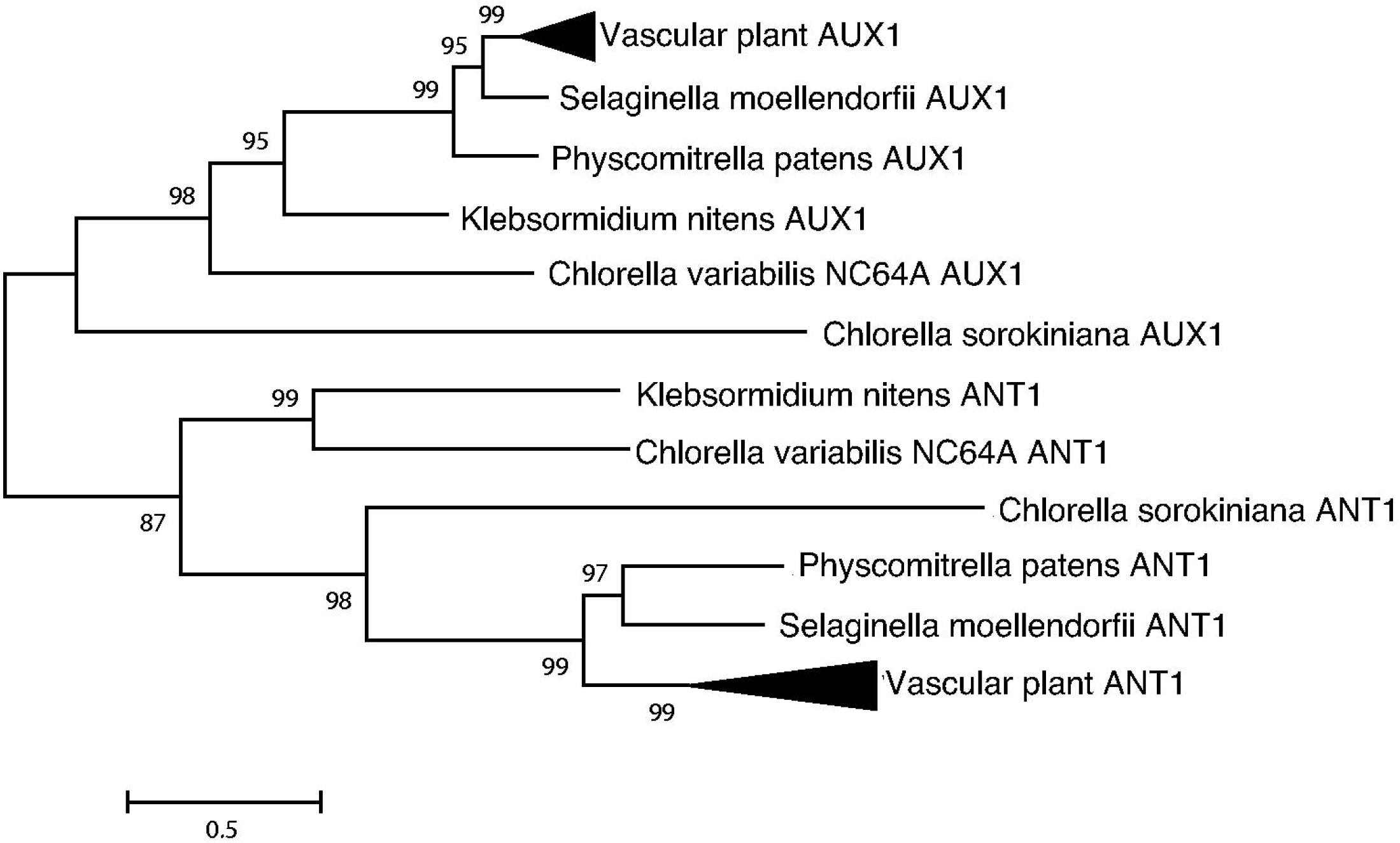
Phylogenetic analysis indicates sequence conservation of AUX1. AUX1, an AAP-like transporter specific for auxin transport, is compared with ANT1, an aromatic amino acid transporter in plants. Putative orthologs and outgroups were identified by reciprocal BLAST searches, aligned with Clustal Omega and analyzed using the LG+I substitution model in MEGA 6.06.

**Figure 4.**
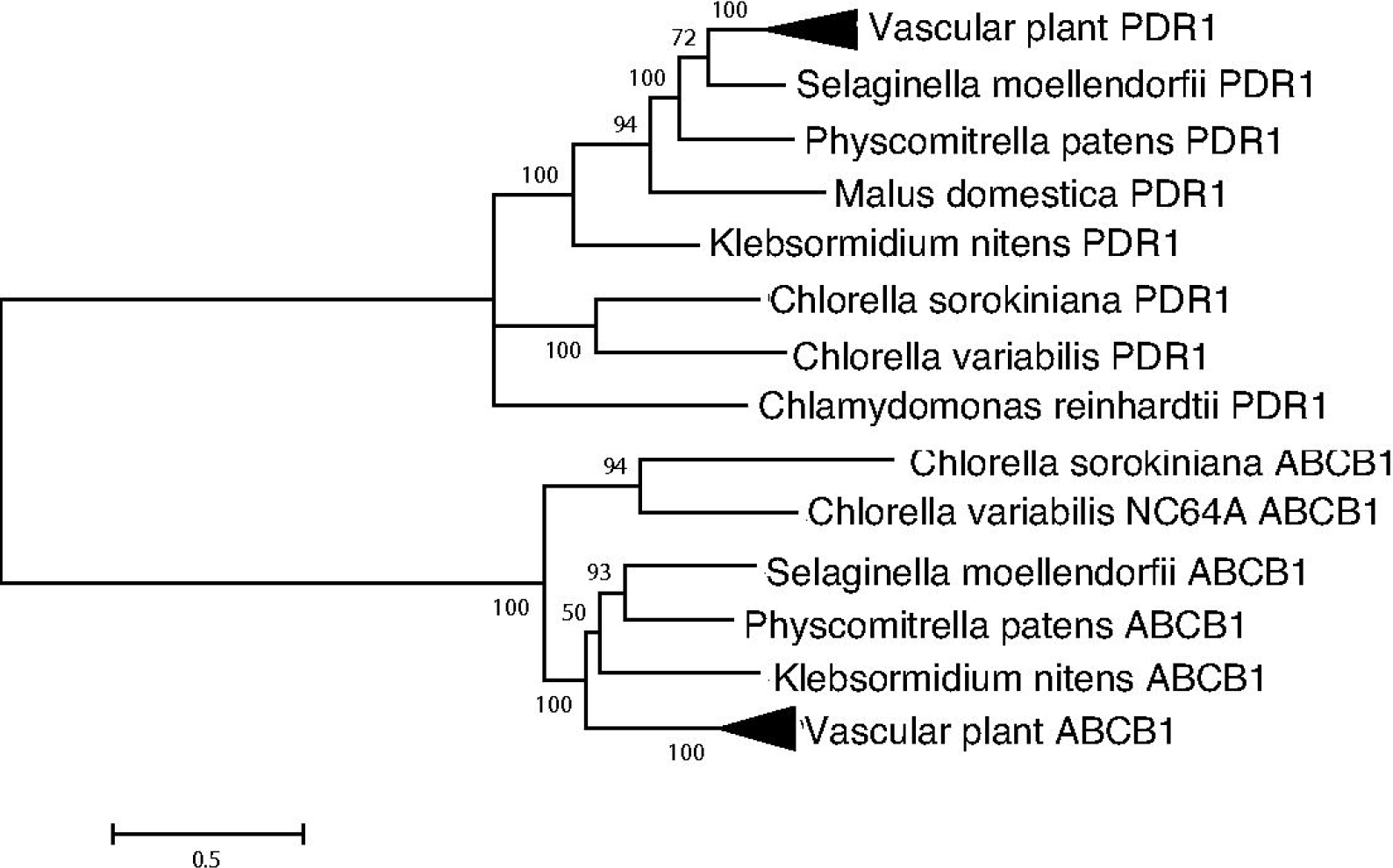
Phylogenetic analysis indicates sequence conservation of ABCB4. An ABCB transporter, is compared with a *p-*coumaric acid transporter which also transports a small organic molecule. Putative orthologs and outgroups were identified by reciprocal BLAST searches, aligned with Clustal Omega and analyzed using the LG+I substitution model in MEGA 6.06.

The redundancy of auxin related genes in the *Arabidopsis* genome suggests a multitude of functions that can be finely modulated and thus, it is possible that the single auxin related genes in *C. sorokiniana* control a range of physiological processes. The presence of PILs-like intracellular orthologs and the absence of PIN orthologs, which mediate extracellular polar auxin transport, are consistent with a unicellular lifestyle. The later appearance of PIN proteins in plants and in algae with differentiated organs is consistent with their participation in activities related to a multicellular lifestyle.

### ABP1 mediated signal transduction

In higher plants, auxin is perceived by at least two types of receptors: the SCF/TIR1/AFB co-receptors, responsible for transcriptional responses to auxin, and ABP1, a cupin domain containing protein whose signaling primarily effects rapid, nontranscriptional auxin responses such as cell expansion and ion fluxes at the plasma membrane [3]. *C. sorokiniana* does not contain orthologs to the SCF/TIR1/AFB co-receptors nor to the auxin responsive transcription factor (ARF). However, it does contain a highly conserved ortholog to ABP1 (Table 1, Figure 5).

**Figure 5.**
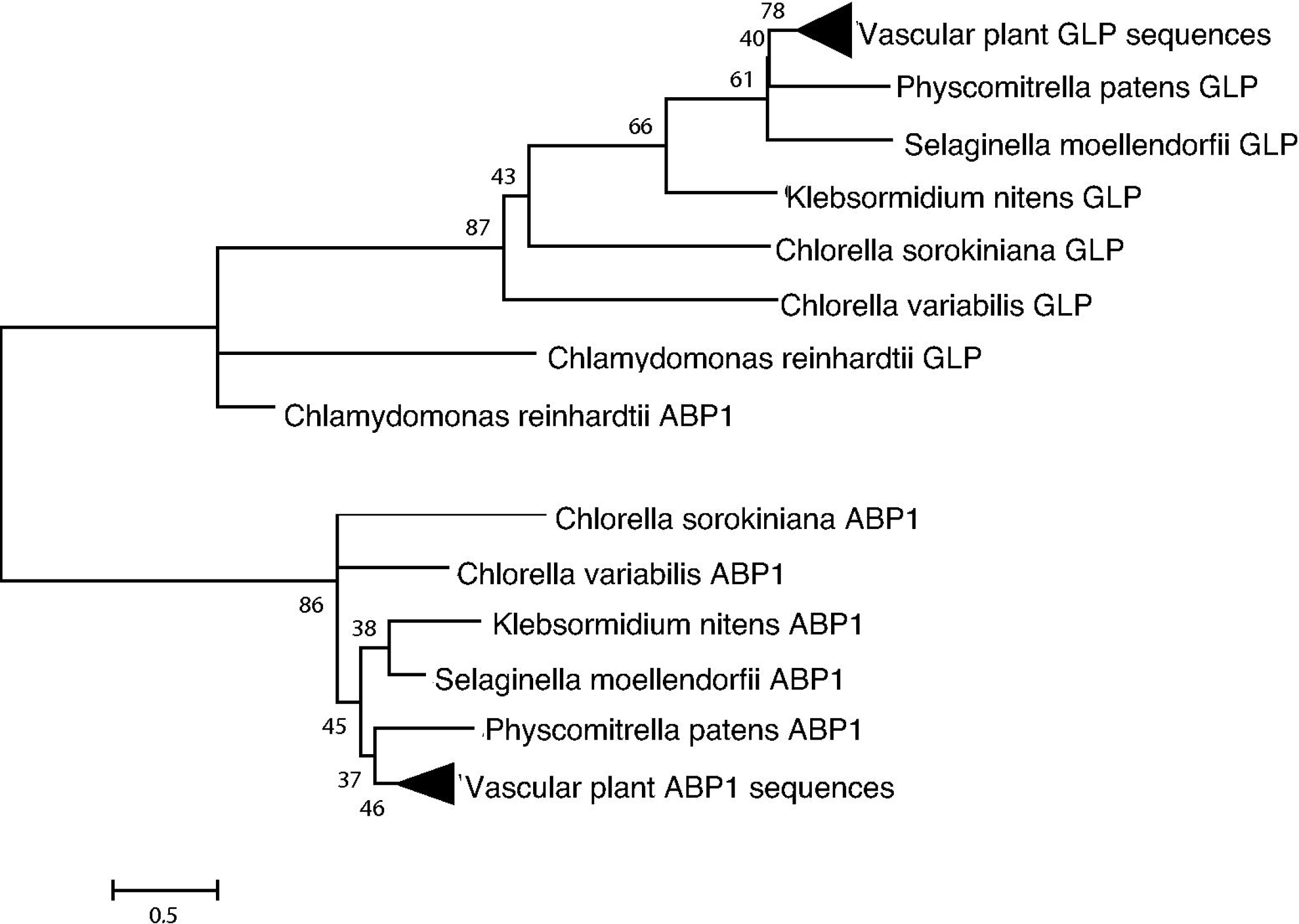
Phylogenetic analysis indicates sequence conservation of ABP1. ABP1, a putative IAA receptor, compared with GLP1, a similarly short (145 AA) cupin domain protein. Putative ABP1 ortholog and outgroups were identified by reciprocal BLAST searches, aligned with Expresso and analyzed using the LG+I substitution model in MEGA 6.06.

ABP1 is a putative auxin receptor first discovered in cell fractions from maize coleoptiles [30]. It was characterized as an auxin receptor in the early 1990s, and the protein structure was solved in 2001. Woo et al. (2001) crystallized ABP1 with and without the synthetic auxin napthaleneacetic acid (NAA) [13]. The structure determined that the auxin binding region features a cluster of hydrophobic amino acids that bind the aromatic ring(s) of auxin, as well as three histidines and a glutamic acid which coordinate a zinc ion which interacts with the carboxylate moiety of IAA. We identify ABP1 as the putative auxin receptor in *Chlorella* species. Structure-based sequence alignment by Expresso [15] followed by pairwise structural alignment by SWISSMODEL [23] (Figure 6) revealed 42% identity and a high degree of structural similarity between *Z. mays* ABP1 and *C. sorokiniana* ABP1.

**Figure 6.**
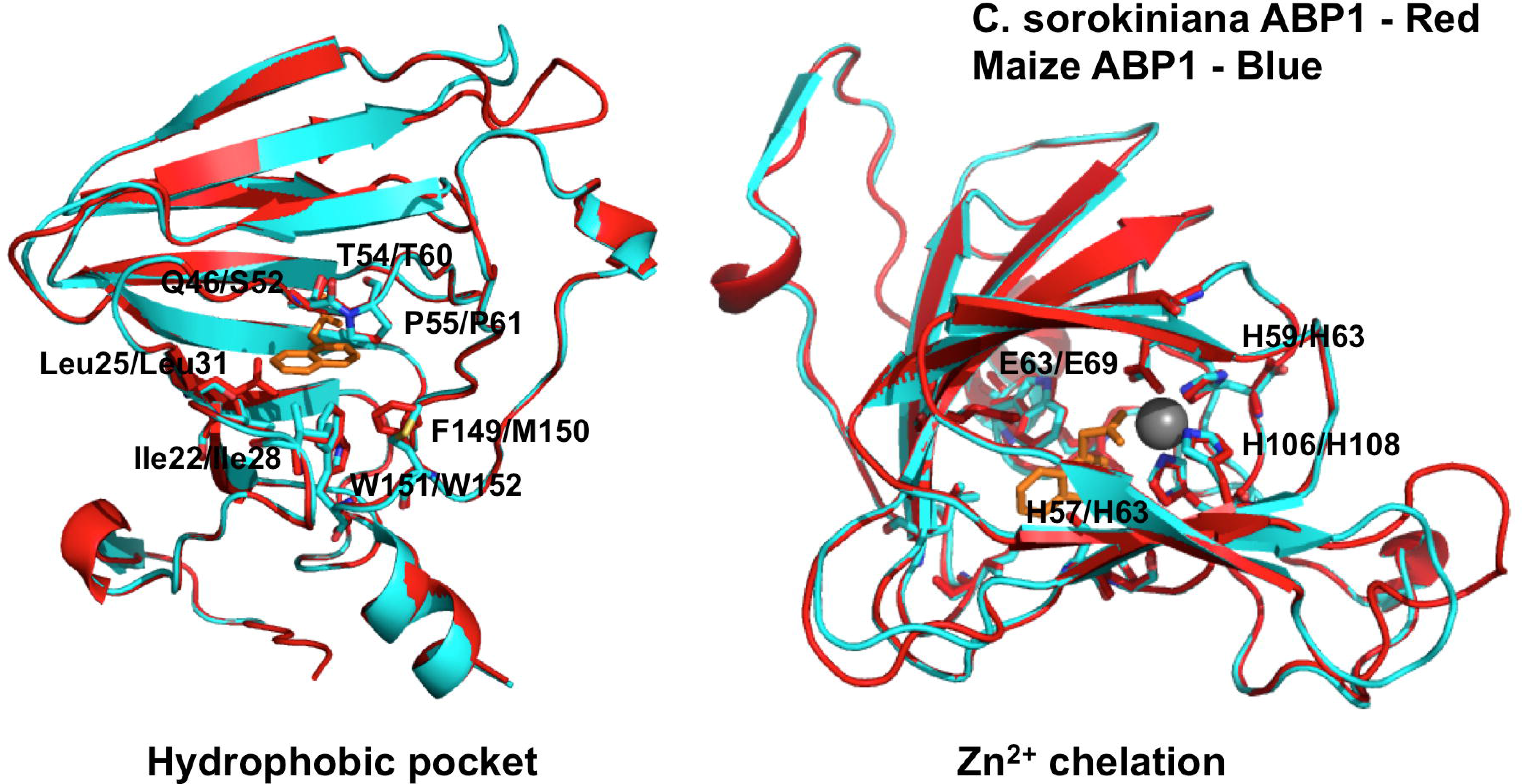
Multiple sequence and structural alignment reveal a high degree of structural conservation between *Z. mays* ABP1 and *C. sorokiniana* ABP1. A SWISSMODEL structural alignment on automated mode aligned CsABP1 with the ZmaABP1 crystal structure co-crystallized with the synthetic auxin naphthalenacetic acid (NAA). The resulting model revealed almost complete conservation of the auxin binding pocket **(A)**, and complete conservation of the histidines and and glutamic acid residues that coordinate the zinc atom **(B)** that stabilizes the carboxylate moiety of IAA/NAA. The red structure represents *Zea mays*, and the blue structure represents *Chlorella sorokiniana*. *Z. mays* residues are indicated first, with *C. sorokiniana* residues following the slash.

The auxin binding pocket is almost completely conserved, except for a phenylalanine to methionine substitution and a glutamic acid to serine substitution. Additionally, the histidines and glutamic acid residues coordinating the zinc atom were completely conserved (Figure 6). This high degree of conservation in key residues suggests a functional role for the ABP1 ortholog in *Chlorella*. In *Arabidopsis*, auxin-bound ABP1 is essential for the activation of Rho-dependent GTPases (ROPs), mediated by plasma membrane associated transmembrane kinases (TMKs) [31]. *C. sorokiniana* encodes both ROP and TMK orthologs for this signaling pathway (Table 1, Figure 1). Additionally, *C. sorokiniana* encodes an ortholog for IBR5, a recently identified TIR-interacting indole butyric acid receptor, but putative functions and interactions for IBR5 in *C. sorokiniana* remain undefined [21].

**Figure 7.**
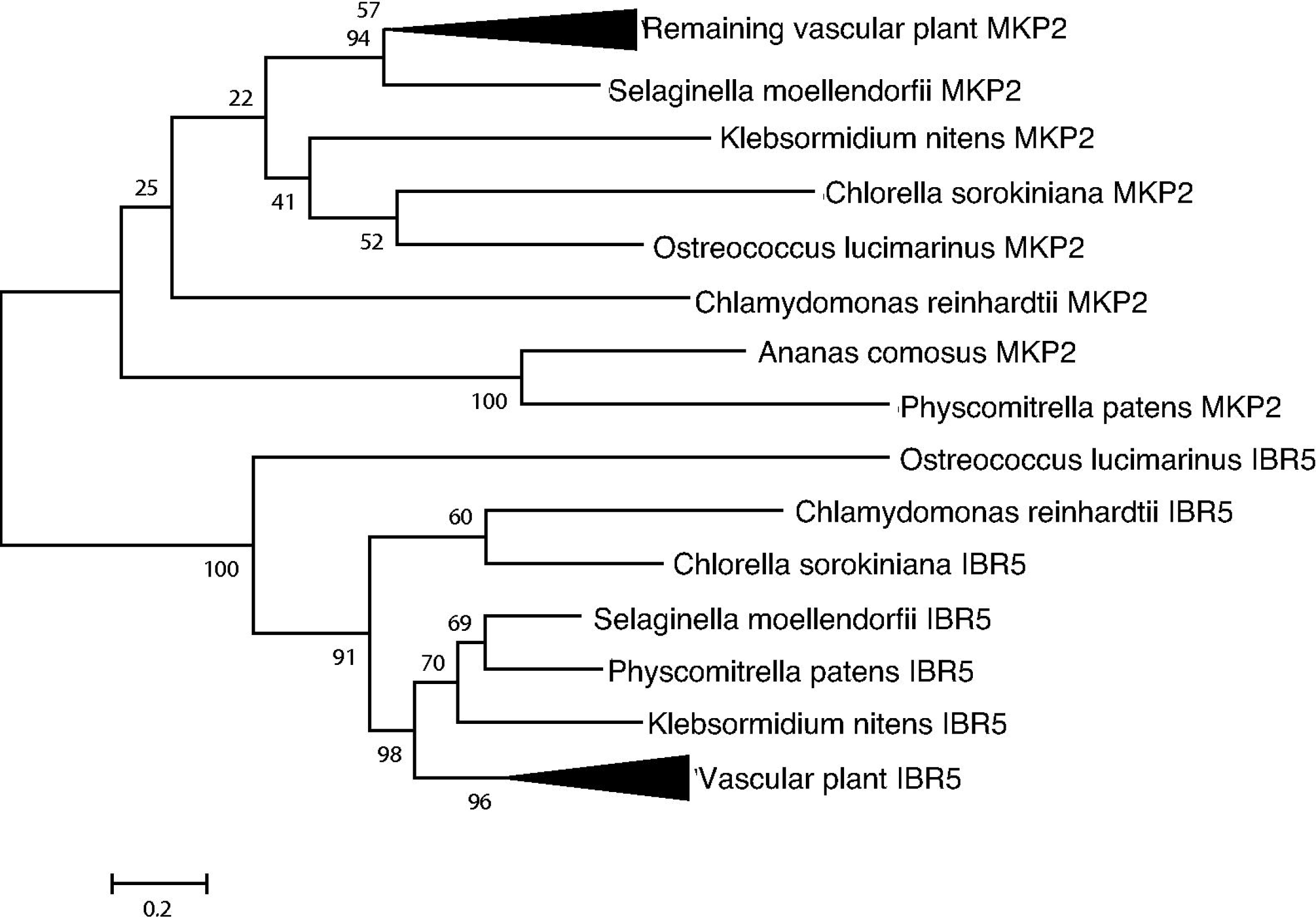
Phylogenetic analysis indicates sequence conservation, suggesting bona fide IAA signaling orthologs in *Chlorella*. IBR5, identified as an indole-3-butyric acid specific receptor, is compared with MKP2, which also contains a MAP kinase phosphatase domain and participates in phytohormone signaling pathways. Putative auxin signaling orthologs and outgroups were identified by reciprocal BLAST searches, aligned with Clustal Omega (except for ABP1, aligned with Expresso) and analyzed using the LG+I substitution model in MEGA 6.06.

The presence of ABP1 and IBR5 and the absence of AUX/IAA signaling components suggests that non-AUX/IAA mediated signaling pathways in plants may have originated in algae, and that their function broadened and diversified as plants colonized land.

### Synthesis and secretion of IAA into the medium

Under the conditions tested, *C. sorokiniana* produces (per 50 ml culture) approximately 2 ng of IAA in the cell pellet and secretes 9 ng into the culture supernatant, resulting in a supernatant concentration of approximately 1 nM (Figure 8).

**Figure 8.**
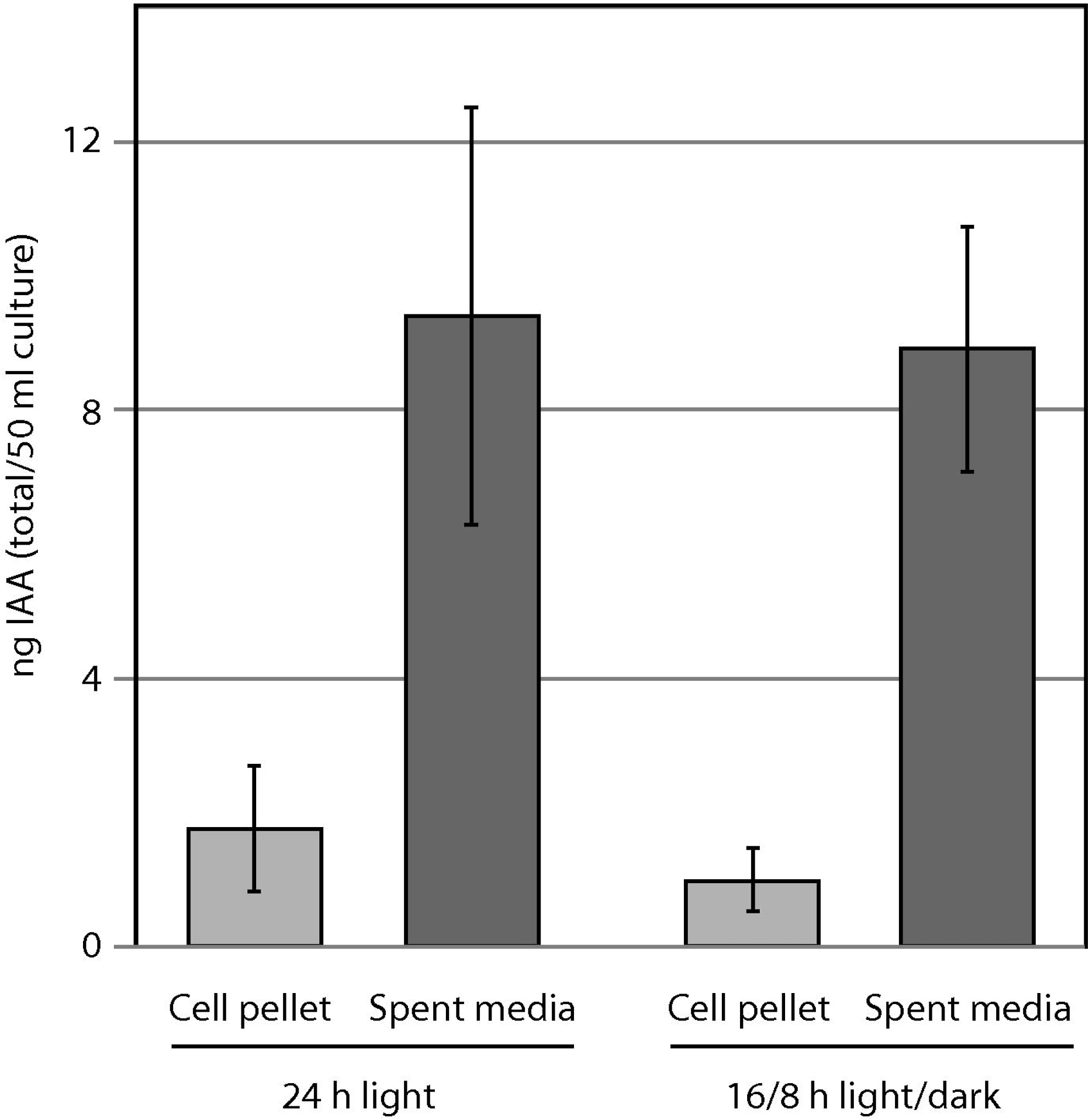
IAA production in *Chlorella sorokiniana*. *Chlorella sorokiniana* produces IAA within the cell pellet and secretes it into the culture medium. For each 50 ml culture, approximately 2 ng is retained within the cell pellet, while 8-9 ng is secreted into the culture medium.

In *Arabidopsis*, the TIR1/AFB receptors perceive IAA at a *K*_d_ of 84 nM [32], and purified *Z. mays* ABP1 has a K_*d*_ of 100 nM for IAA [33]. However, *C. sorokiniana* has been reported to grow as a biofilm in biofuel and waste remediation biotechnology, and we have observed biofilm growth in glass Erlenmeyer growth flasks, in agreement with prior reports [34,35]. In a biofilm context, the concentration of IAA in the extracellular matrix between cells would be at least 2 orders of magnitude higher than that observed in our bulk planktonic cultures, placing the in situ IAA concentration well within the physiologically relevant range when compared to IAA concentrations in higher plant tissues.

## Conclusions

The conservation of many auxin synthesis, transport, and signaling related orthologs in *Chlorella sorokiniana* and other unicellular algae demonstrates that a core set of genes required for auxin signaling were present in unicellular chlorophytes, and the production and secretion of IAA into the medium shows that the synthesis and secretion pathways are operative. However, it must be emphasized that the specific function(s) of IAA in algae are currently poorly defined, and while our work provides a foundation upon which new studies can be built, we can currently only speculate as to the functions of auxin synthesis, secretion, and signaling in unicellular chlorophytes.

A number of possible functions for auxin as an inter-or intracellular signaling molecule are conceivable. First, given our demonstration that IAA is a secreted molecule, one possibility is that IAA signaling evolved as a population-density dependent mediator of quorum sensing in algae, analogous to bacterial and fungal quorum sensing molecules. Another possibility is that IAA acts as a regulator of biofilm initiation and development, whereby the molecule is constitutively secreted and maintained at a high concentration in biofilms, and at a much lower concentration when cells are not adhered to a substrate. Still another possibility is that IAA acts as an interspecies signaling molecule, allowing algae to communicate with members of mixed biofilm communities.

From the perspective of the evolution of multicellularity and pattern formation in the green plant lineage, this work provides compelling evidence that the molecular machinery necessary for auxin signaling was present in the unicellular ancestors of higher plants, and was thus available for elaboration and specification of auxin signaling pathways during the evolution of land plants.

## Acknowledgments

This work was funded by Nebraska Center for Energy Sciences Research (NCESR) and the National Science Foundation (EPS-1004094).

